# Loss of HIF1α signaling drives oxidative stress and expansion of smooth muscle cells in murine atherosclerosis

**DOI:** 10.64898/2026.06.26.734925

**Authors:** Raúl Izquierdo-Serrano, Diana Sharysh, Vanessa Cumbicus, Pablo Hernansanz-Agustín, Judith C. Sluimer, Silvia Martin-Puig, Laura Carramolino, Daniel Morales-Cano, Jacob F Bentzon

**Affiliations:** Centro Nacional de Investigaciones Cardiovasculares Carlos III, Madrid Spain; Atherosclerosis Research Unit, Department of Clinical Medicine, Aarhus University, Aarhus Denmark; Departamento de Neurobiología Molecular, Celular y del Desarrollo, Centro de Neurociencias Cajal (CNC), Consejo Superior de Investigaciones Científicas (CSIC), Madrid, Spain; Department of Pathology, Cardiovascular Research Institute Maastricht, Maastricht University Medical Center, Maastricht, the Netherlands; Department of Medicine 2 (Nephrology, Clinical Immunology, Rheumatology, Hypertension), RWTH Aachen University, Medical Faculty, Aachen, Germany; BHF centre for Cardiovascular Sciences, University of Edinburgh, Edinburgh, UK; Instituto de Investigaciones Biomédicas Sols-Morreale, (IIBM), Consejo Superior de Investigaciones Científicas & Universidad Autónoma de Madrid (CSIC-UAM), Madrid, Spain; Departamento de Farmacología, Facultad de Medicina, Universidad Complutense, Madrid, Spain; Steno Diabetes Center Aarhus, Aarhus University Hospital, Aarhus, Denmark

**Author notes:** **Corresponding authors:** Raúl Izquierdo Serrano, Centro Nacional de Investigaciones Cardiovasculares C/Melchor Fernandéz Almagro, 3, 28029 Madrid, Spain, Jacob F. Bentzon, MD, PhD, Department of Clinical Medicine, Aarhus University, Palle Juul-Jensens Boulevard 11, 8200 Aarhus N, Denmark.

**Keywords:** atherosclerosis, smooth muscle cells, hypoxia, hypoxia-inducible factor 1 alpha, oxidative stress

## Abstract

**Background:** Hypoxia develops within growing atherosclerotic lesions, inducing nuclear translocation of hypoxia-inducible factor-1α (HIF1α) and metabolic reprogramming. Its role in plaque macrophages and endothelial cells has been studied, but the hypoxic plaque interior is dominated by smooth muscle cell (SMC)-derived cells, for which the role of hypoxia signaling remains unclear. Here, we investigated how loss of *Hif1a* in SMC lineage cells impacts plaque progression and cell phenotype in murine atherosclerosis.

**Methods:** Atherosclerosis was induced in mice with inducible SMC-specific deletion of *Hif1a* (*Hif1a^SMC-KO^*) and lineage tracing of SMC-derived plaque cells. Plaque size, necrotic core size, calcification, and SMC-derived cell phenotypes were quantified in aortic root sections and gene expression changes mapped by single-cell RNA sequencing. In parallel, a cultured SMC line with or without siRNA-mediated *Hif1a* knockdown was exposed to hypoxia for assessments of mitochondrial function and reactive oxygen species production.

**Results:** *Hif1a^SMC-KO^* mice developed larger plaques, with expanded necrotic cores and increased calcification, compared with littermate controls. SMC-derived plaque cells were more abundant with a higher fraction of *Col2a1*^+^ chondromyocytes, and showed elevated markers of proliferation and apoptosis, whereas macrophage and endothelial cell numbers were unaffected. Single-cell RNA sequencing analysis revealed strong dysregulation of mitochondrial genes, including electron transport chain transcripts, along with upregulation of protein folding, proteasome, and oxidative stress response pathways. In cultured SMCs subjected to hypoxia, *Hif1a* silencing increased cell counts, aggravated mitochondrial proton leak, and led to the accumulation of depolarized, reactive oxygen species-generating mitochondria. Further analysis of SMC-derived cells in plaques from *Hif1a^SMC-KO^* mice confirmed increased oxidative stress by 8OHdG staining.

**Conclusions:** HIF1α maintains mitochondrial function and restrains oxidative stress in SMC-derived plaque cells in murine atherosclerosis. Its chronic loss destabilizes redox homeostasis and promotes maladaptive SMC responses, leading to SMC-driven plaque expansion, necrosis, and calcification.

## Introduction

Atherosclerotic plaque expansion in humans, pigs, rabbits, and mice increases the diffusion distance for oxygen, producing hypoxia in the plaque interior.^1–4^ The low oxygen tension may, in turn, accelerate atherosclerosis through multiple processes, creating self-perpetuating cycles. Previous studies have focused on macrophages and endothelial cells, showing that plaque hypoxia impairs lipid efflux, promotes monocyte recruitment, and alters macrophage polarization;^5,6^ however, its effects on other prevalent cell types in atherosclerosis remain poorly understood.

SMC-derived plaque cells constitute the largest cell population in advanced murine atherosclerosis,^7^ encompassing cap SMCs, fibroblast-like cells (fibromyocytes), and chondrocyte-like cells (chondromyocytes).^8^ These phenotypes exhibit a distinct spatial pattern, with cap SMCs localized near the endothelium and fibromyocytes and chondromyocytes concentrated in the plaque core.^9,10^ This organization likely exposes SMC-derived cell types to different oxygen levels, yet the role of hypoxia signaling in SMC-derived cell growth and phenotypic modulation is unknown.

A central mediator of the cellular hypoxia response is hypoxia-inducible factor 1α (HIF1α). Under normoxia, HIF1α is hydroxylated at specific proline residues by prolyl hydroxylases, enabling proteasomal degradation by the von Hippel–Lindau tumor suppressor (VHL). In hypoxia, prolyl hydroxylase activity is inhibited, allowing HIF1α to escape degradation and dimerize with HIF1β to form the HIF1 heterodimer, which translocates to the nucleus to activate hypoxia-responsive genes via binding to hypoxia response elements (HREs).^11^

In addition to regulating gene expression through HIF1 complexes, hypoxia affects intracellular redox homeostasis,^12^ counterintuitively increasing mitochondrial reactive oxygen species (ROS) production.^13^ This occurs due to partial inhibition of the electron transport chain, which causes the accumulation of semi-reduced intermediates at complex III that donate electrons to molecular oxygen, generating superoxide and other ROS.^14^ The elevated ROS level further stabilizes HIF1α by inhibiting prolyl hydroxylases, thereby amplifying HIF1α accumulation and transcriptional activity.^15^ As HIF1α target genes encode antioxidant defenses, this establishes a feedback loop that regulates ROS levels to maintain them within a physiological range.^16,17^ ROS can thus act both as signaling molecules and produce cellular damage, depending on concentration, location, and cellular context. The role of this hypoxia-ROS-HIF1α axis in SMC-derived cells is largely unexplored.

In this study, we combined SMC-specific *Hif1a* deletion, lineage tracing, *in vitro* assays, and single-cell RNA sequencing to define the role of hypoxia signaling in SMC-derived plaque cells. We find that loss of HIF1α increases the abundance of ROS-producing mitochondria, driving oxidative stress and yielding larger atherosclerotic plaques enriched in SMC-derived cells, with expanded necrotic cores and calcified regions.

## Methods

The raw data that support the findings of this study are available from the corresponding authors on reasonable request. ScRNA-seq data have been deposited in ArrayExpress with accession number (E-MTAB-17142).

### Animals

To obtain mice in which HIF1α signaling can be disrupted in lineage-traced SMCs, we intercrossed *Myh11-CreER^T^*^2^ mice (B6.FVB-Tg[Myh11-cre/ERT2]1Soff/J, The Jackson Laboratory), expressing tamoxifen-inducible Cre recombinase (CreER^T2^) under the SMC-specific *Myh11* promoter; tdTomato (tdT) reporter mice (B6.Cg-Gt(ROSA)26Sortm14(CAG-tdTomato)Hze/J), where Cre activity activates a tdT-encoding transgene; and *Hif1a* floxed mice (B6.129-*Hif1a*^tm3Rsjo^/J), in which Cre deletes exon 2 encoding the dimerization domain.^18^ Lines were extensively backcrossed to the C57BL/6J genetic background. Mice compared in experiments were littermates produced by matings of *Myh11-CreER^T^*^2^-tdT^tg/tg^-*Hif1a^flx/wt^*and *Hif1a^flx/wt^* mice, housed together, subjected to the same procedures, and differing only in genotype. Only male offspring were studied since the *Myh11-CreER^T^*^2^ transgene is inserted on the Y chromosome.

Mice were group-housed under specific pathogen-free conditions with a 12-h light–dark cycle at 23 ± 1°C and 50 ± 5% humidity with *ad libitum* access to water and food (SAFE D184, Scientific Diets, when not on high-fat diet) at the Centro Nacional de Investigaciones Cardiovasculares Carlos III (CNIC). Experiments were approved by the CNIC Ethical Committee for Animal Welfare and by the Comunidad de Madrid (PROEX 020.8/21).

### Induction and atherosclerosis development

Mice were injected intraperitoneally with tamoxifen (2 mg) dissolved in corn oil on five consecutive days starting at 6 weeks of age. A subset of mice was maintained until 30 weeks of age to provide non-atherosclerotic controls for evaluation of *Hif1a* deletion efficiency. Other mice received a single tail vein injection of recombinant adeno-associated virus encoding liver-expressed proprotein convertase subtilisin/kexin type 9 (rAAV8-PCSK9) particles at 9 weeks of age (1×10^11^ vector genomes, produced in the CNIC viral vector core facility), followed by feeding with a high-fat diet (SSNIFF-S9167-E011) to induce atherosclerosis. Plasma cholesterol was measured in duplicate in serial blood samples using an enzymatic cholesterol assay (CH201, Randox Reagents). After 21 weeks of atherogenesis (30 weeks of age), mice were anesthetized (250 mg/kg pentobarbital and 20 mg/kg lidocaine). Blood was collected via puncture of the right ventricle and mice were euthanized by exsanguination. The heart was stopped in diastole by perfusion of 50 mM KCl and residual blood was flushed from the circulation with cold phosphate-buffered saline (PBS). Tissues were immersion fixed in 4% phosphate-buffered formaldehyde for 24 hours stored in PBS at 4°C until embedding. In a subset of atherosclerotic mice, aortic arches were harvested before immersion fixation and processed for scRNA-sequencing analysis as described in detail in Supplemental Methods.

### Histological analysis

Hearts were extracted, cryoprotected in 25% and 50% sucrose solutions and frozen in Optimal Cutting Temperature compound (OCT). Serial 5µm cross-sections were collected at the level of the aortic valve commissures (0 µm level) and at levels 100 µm and 200 µm distal to this reference point. Histological analyses (plaque size, necrotic core size) were performed on all available animals and all levels. Immunofluorescence staining was done on sections at the 100 µm level in a subset of animals as specified in figure legends.

Overall plaque morphology and size were determined by averaging measurements from four aortic root sections per mouse, whereas other data represent measurements from a single section per mouse. Necrotic cores were defined as large, cell-free regions indicative of advanced lesion development and cell death. Masson’s trichrome staining was used to identify acellular areas larger than 1000 μm²-excluding isolated cholesterol crystals-which were quantified as indicators of necrotic core formation. Alcian blue staining was performed to detect acidic polysaccharides such as glycosaminoglycans. Lipid content within lesions was assessed using Oil Red O staining, which labels neutral triglycerides and lipids. In addition, Alizarin Red staining was used to evaluate calcium deposits within the aortic root lesions.

### Statistical analysis

Two-group comparisons were performed with unpaired, two-tailed Student’s t-test after confirming normal distribution. When the outcome consisted of matched averages from the same experiment repeated n times, a paired, two-tailed Student’s t-test was applied to account for intra-experiment variability. Multi-group datasets were analyzed by one-way ANOVA or, when paired measures were present, by RM one-way ANOVA with Greenhouse–Geisser correction, both followed by Tukey’s multiple-comparison test. Experiments involving two independent factors (e.g., genotype × treatment) were evaluated with two-way ANOVA followed by Tukey’s post-hoc test. Calcification status was treated as a binary trait and compared between genotypes with Fisher’s exact test. Statistical significance was set at P < 0.05. Calculations and plotting were performed in GraphPad Prism 10 (GraphPad Software), except for single-cell RNA-seq gene-set score comparisons, which were carried out separately in R (v 4.4.1) using the Wilcoxon rank-sum test with continuity correction from the base ‘stats’ package.

No exclusions of data were made, but a few data points are missing for technical reasons, e.g. folded tissue sections or failed blood sampling. The number of replicates in each analysis is specified in figure legends.

RNA extraction, quantitative PCR, immunofluorescence, in situ hybridization, scRNA-seq analysis, SMC culture, and flow cytometry is described in the Supplementary material. Details for antibodies are provided in **Table S1-S2**.

## Results

### Loss of functional HIF1α in SMC-lineage cells exacerbates atherosclerosis

To examine the role of HIF1α signaling in SMC-lineage cells, we generated mice with SMC-specific deletion of *Hif1a* exon 2 (*Hif1a*^SMC-KO^) using a tamoxifen-inducible Myh11-CreER^T2^ driver at 6 weeks of age (**Fig. 1A**).^18^ Deletion of *Hif1a* exon 2 yields a truncated HIF1α protein that cannot bind HIF1β and translocate to the nucleus to exert transcriptional activity, while still undergoing VHL-mediated degradation, thereby avoiding imbalances in other VHL substrates.^19^ Littermate and co-housed mice with wildtype *Hif1a* alleles (*Hif1a*^WT^) served as controls, and all mice carried a Cre-activable tdTomato reporter enabling SMC lineage tracing. Efficient and persistent deletion was confirmed by quantitative PCR of the aortic media, which showed ≈80% reduction in *Hif1a* exon 2 expression in *Hif1a*^SMC-KO^ (n=8) compared with *Hif1a*^WT^ (n=10) mice at 30 weeks of age (**Fig. 1B**).

**Figure 1.**
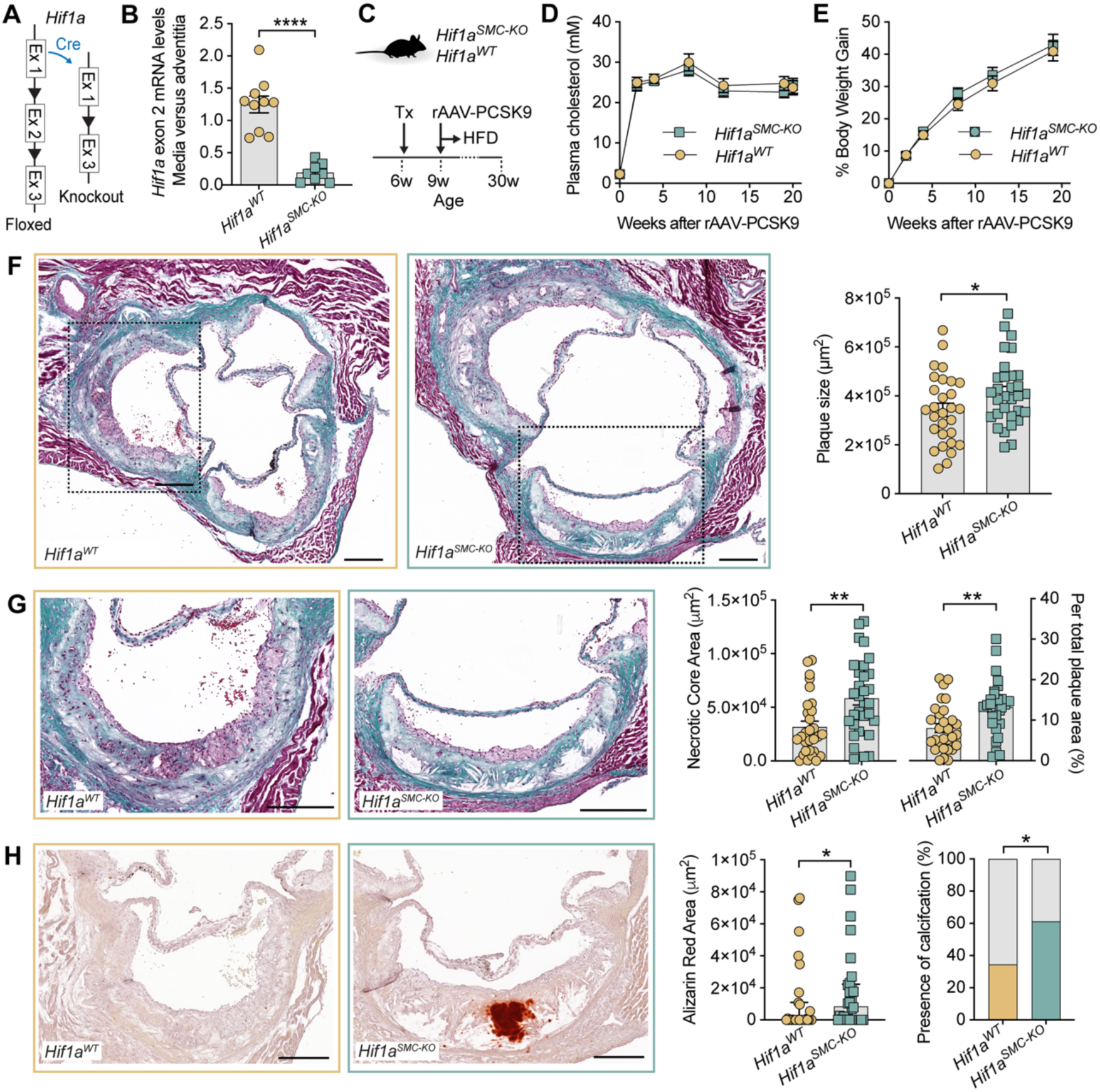
*Hif1a* deletion in SMC-lineage cells exacerbates atherosclerosis. **A,** Schematic representation of tamoxifen-inducible *Hif1a* recombination. **B,** Quantitative RT-qPCR analysis of *Hif1a* exon 2 expression in aortic media, normalized to adjacent adventitia, in *Hif1a*^WT^ and *Hif1a*^SMC-KO^ mice. Samples were obtained at 30 weeks of age. **C,** Experimental design of the atherosclerosis experiment. Tx, tamoxifen. HFD, high-fat diet. **D-E,** Evolution of plasma cholesterol levels and body weight over time in *Hif1a*^WT^ and *Hif1a*^SMC-KO^ mice. **F,** Representative images of aortic root sections stained with Mallory’s Trichrome, along with quantification of plaque size. **G,** Magnified plaque views and quantification of necrotic core size (defined as acellular regions) as absolute values (left) and normalized to plaque area (right). **H,** Representative images and quantification of Alizarin Red-stained plaque calcification. The panels display absolute values (left) and a contingency analysis of the presence of any plaque calcification (right). Bars, mean ± SEM (D-G) or median and interquartile range (H). Scale bars, 200 µm. Data were obtained from 10 *Hif1a*^WT^ and 8 *Hif1a*^SMC-KO^ mice in B and 28-30 *Hif1a*^WT^ and 31-32 *Hif1a*^SMC-KO^ mice in C-H. Continuous data was compared between groups by unpaired two-tailed t-tests (D-G, using the area under the curve in D-E) or Mann Whitney test (H). Contingency analysis was performed by Fisher’s exact test (H). \**P*<0.05, **P<0.01.

Atherosclerosis was induced in 9-week-old *Hif1a*^SMC-KO^ (n=31) and *Hif1a*^WT^ (n=29) mice by the injection of rAAV-PCSK9 followed by feeding a high-fat diet until 30 weeks of age (**Fig. 1C**). No significant differences in plasma lipids or body weights were observed during the study (**Fig. 1D-E**), but *Hif1a*^SMC-KO^ mice developed larger plaques in the aortic root than *Hif1a*^WT^ mice, with increased necrotic cores and enhanced calcification (**Fig. 1F-H**). Plaque proteoglycan-rich extracellular matrix (Alcian blue-stained area) and lipids (Oil Red O-stained area) were also measured but showed no significant changes (**Figure S1**).

### Hif1a^SMC-KO^ drives SMC-derived cell accumulation, proliferation, and apoptosis

To determine whether the exacerbated disease involved changes in the cellular composition of plaques, we stained sections for CD31 and CD68 and used the endogenous tdT lineage tag to identify SMC-derived cells (**Fig. 2A**). Absolute numbers of SMC-derived (tdT^+^) cells were significantly increased in *Hif1a^SMC-KO^* relative to *Hif1a*^WT^ plaques, whereas numbers of macrophages (CD68^+^) or endothelial cells (CD31^+^) remained unchanged (**Fig. 2B**). To assess whether expansion of the SMC-derived population reflected altered proliferation or cell death, we stained for Ki67 (a marker of cells in active phases of the cell cycle) and TUNEL (a marker of apoptotic DNA fragmentation). The numbers of tdT⁺ Ki67⁺ (**Fig. 2C-D**) and tdT⁺ TUNEL⁺ (**Fig. 2E-F**) cells were both increased in plaques of *Hif1a*^SMC-KO^ compared with *Hif1a*^WT^ mice, indicating elevated SMC-derived cell proliferation as well as increased SMC-derived cell death, consistent with the observed combination of an expanded SMC-derived population and a larger necrotic core.

**Figure 2.**
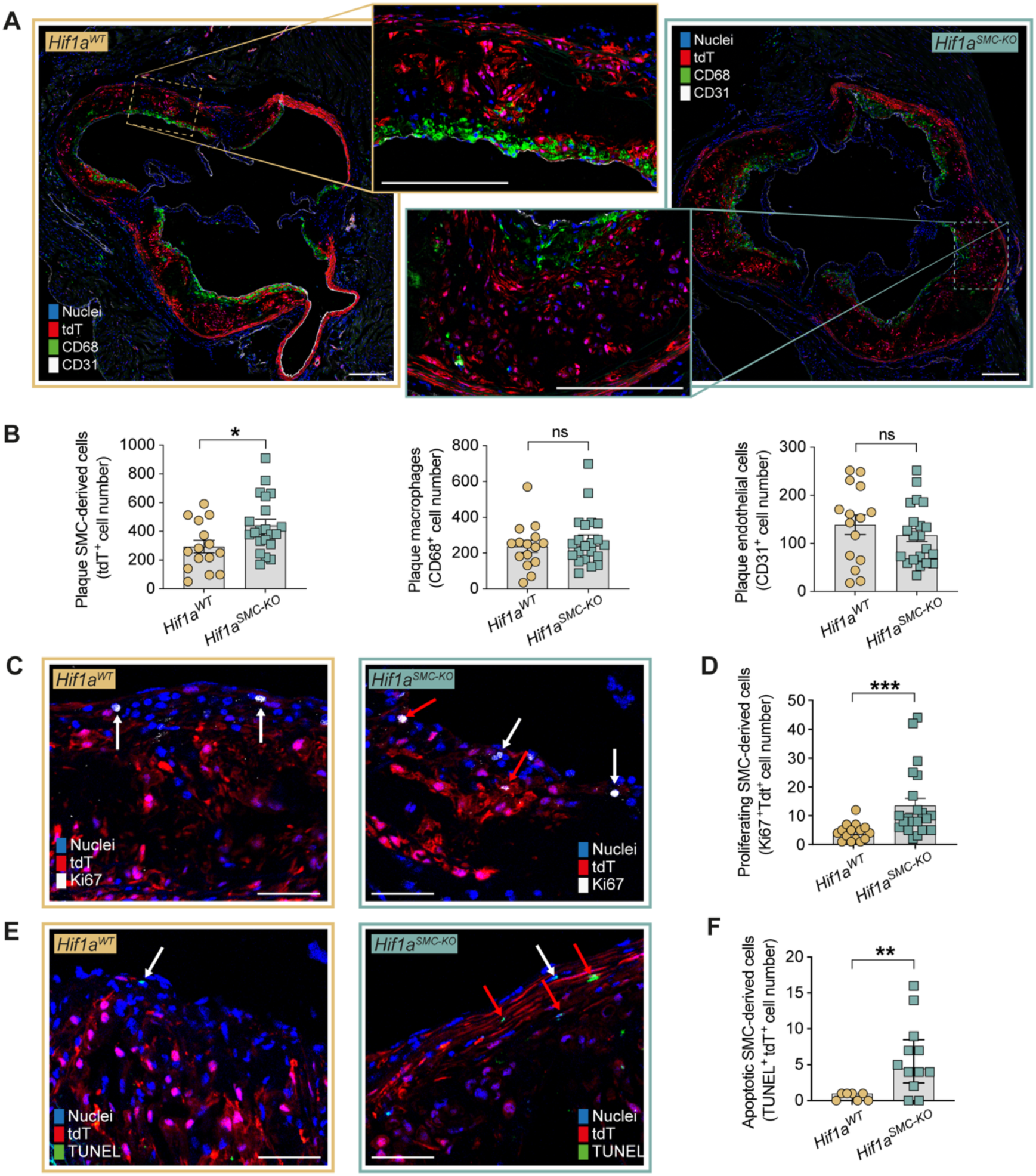
*Hif1a* deletion increases the abundance, proliferation and apoptosis of SMC-derived cells. **A,** Representative immunostaining of aortic root sections for CD68 (macrophages) and CD31 (endothelial cells), with tdTomato (tdT) marking SMC-derived cells. **B,** Quantification of the indicated cell populations. **C,D** Representative Ki67 staining together with tdT and quantification of proliferating SMC-derived cells (Ki67^+^ tdT^+^). **E,F** Representative TUNEL staining together with tdT and quantification of apoptotic SMC-derived cells (TUNEL ^+^ tdT ^+^). Bars, mean ± SEM (B,D) or median and interquartile range (F). Scale bars, 200 µm (A) or 50 µm (C,E). Data were obtained from 15 *Hif1a*^WT^ and 21 *Hif1a*^SMC-KO^ mice in B and D, and from 7 *Hif1a*^WT^ and 12 *Hif1a*^SMC-KO^ mice in F. Groups were compared by unpaired t-tests (B), unpaired t-test using log-transformed data (D) or Mann-Whitney test (F). \**P*<0.05, \*\**P*<0.01, \*\*\**P*<0.001.

### Hif1a^SMC-KO^ plaques are enriched in Col2a1-expressing chondromyocytes

Next, we assessed whether the expansion of SMC-derived cells involved all or specific subtypes. Using ACTA2 and SOX9 as broad markers to define cap SMCs (ACTA2^+^ tdT^+^), fibromyocytes (ACTA2^−^ SOX9^−^ tdT^+^), and chondromyocytes (SOX9^+^ tdT^+^) as previously described,^20^ we observed an increase across all subsets in *Hif1a^SMC-KO^* versus *Hif1a^WT^* plaques, reaching significance for fibromyocytes and chondromyocytes, with the proportional representation of all subsets among tdT⁺ cells remaining unchanged (**Fig. 3A-B**). To further examine the chondromyocyte population, we performed RNAscope for *Col2a1,* which encodes the major cartilage collagen and serves as a more specific marker for cells with a chondrocyte-like phenotype. *Col2a1* transcripts were detected in tdT⁺ cells localized deep within lesions and the fraction of *Col2a1*⁺ tdT⁺ cells among all tdT⁺ cells was significantly increased in *Hif1a^SMC-KO^* compared with *Hif1a^WT^* plaques (**Fig. 3C-D**). Together, these findings indicate that loss of HIF1 signaling leads to an overall expansion of all types of SMC-derived cells with the strongest effect for SMC-derived cells with a mature chondromyocyte phenotype.

**Figure 3.**
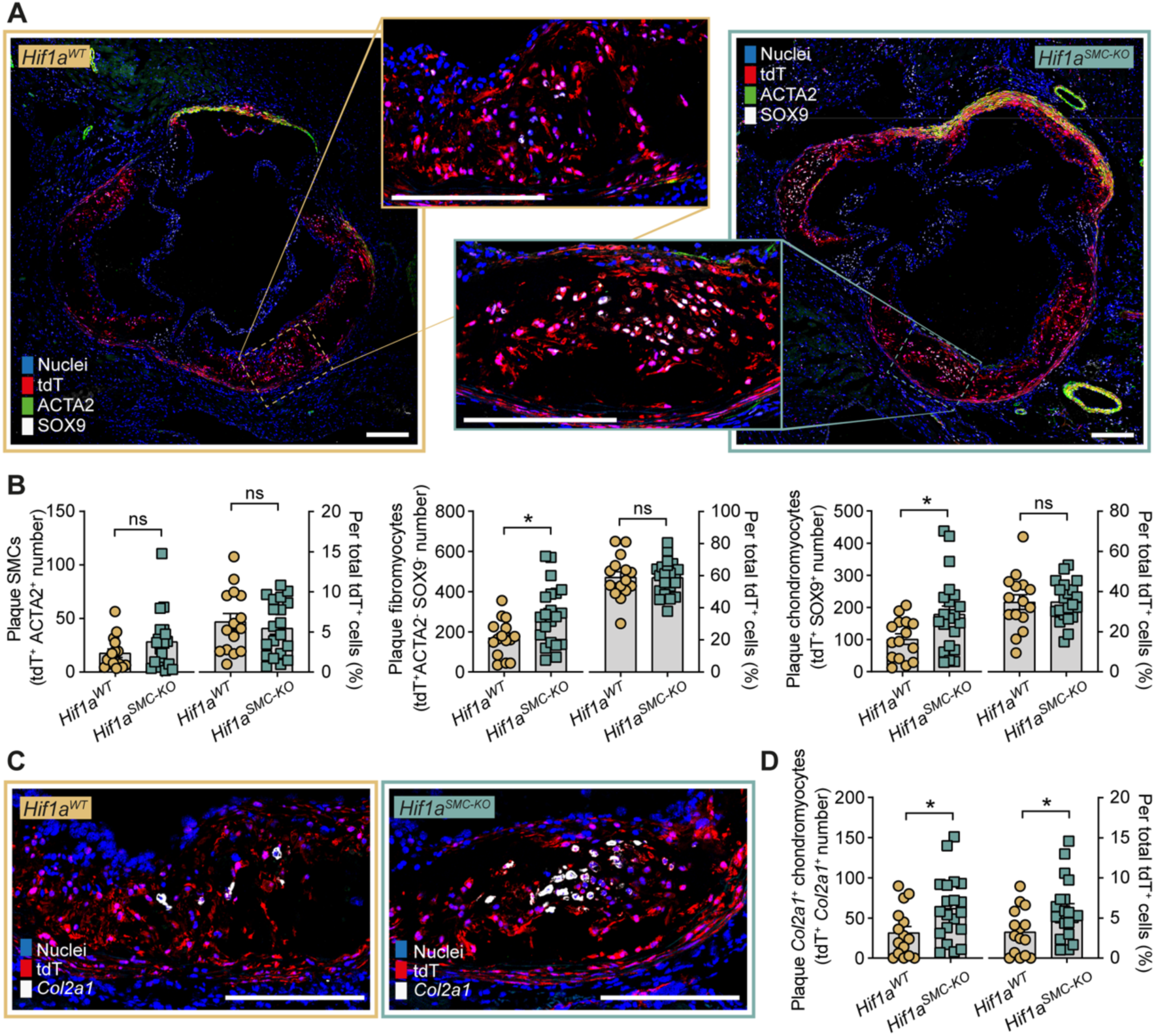
Loss of *Hif1a* skews the SMC-derived cell population towards mature chondromyocytes. **A,** Representative immunofluorescence of ACTA2 and SOX9 in aortic root sections, with tdT marking SMC-derived cells. This marker combination identifies plaque SMCs (ACTA2⁺ tdT⁺), fibromyocytes (ACTA2⁻ SOX9⁻ tdT⁺), and chondromyocytes (SOX9⁺ tdT⁺). **B,** Quantification of plaque SMC-derived cell types. The absolute number of nucleated cell profiles with the indicated marker expression (left) and as a percentage of all tdT+ cells (right) are shown. Triple-positive cells are exceedingly rare (not shown). **C,** Representative RNAscope in situ hybridization for *Col2a1* in aortic root sections, with tdT marking SMC-derived cells, identifying chondromyocytes cells with a more mature *Col2a1*-expressing phenotype. **D,** Quantification of plaque *Col2a1+* tdT+ cells. The absolute number of nucleated *Col2a1*+ cell profiles (left) and as a percentage of all tdT+ cells (right) are shown. Bars, mean ± SEM. Scale bars, 200µm. Data were obtained from 15 *Hif1a*^WT^ and 20-21 *Hif1a^SMC-KO^* mice. Groups were compared by unpaired t-tests. \**P*<0.05.

### Gene expression changes indicate alterations in mitochondria and stress responses

Single-cell RNA sequencing (scRNA-seq) was performed on pooled aortic arches from a subset of the atherosclerotic *Hif1a*^WT^ (n=6) and *Hif1a* ^SMC-KO^ (n=8) mice. To enrich for plaque cells, the adventitia was removed before analysis, but media was retained. After standard quality control filtering and removal of endothelial and immune cells, gene expression profiles were obtained for 2742 and 3466 SMC-derived cells from *Hif1a*^WT^ and *Hif1a*^SMC-KO^ mice, respectively. Dimensionality reduction and clustering at a moderate resolution separated the SMC-derived cells into five subclusters (**Fig. 4A**), consistent with previous reports,^20^ and these were named using a nomenclature adapted from Kim et al.^21^ The following clusters were annotated: SMCs with high contractile gene expression (e.g. *Acta2*, *Myh11*, *Cnn1)* of which some (denoted SMC*) had co-expression of a set of stress-induced genes (e.g. *Fos*, *Irf1*), which may be a tissue-dissociation artefact;^22^ *Rgs5*+ SMCs with high *Rgs5* expression and intermediate levels of contractile gene expression, previously shown to be located both in the fibrous cap and scattered among medial SMCs;^20^ fibromyocytes with low contractile gene expression, expression of extracellular matrix genes (e.g. *Lum*), and markers of inflammatory signaling (e.g. *C3, Vcam1*);^23^ and chondromyocytes with low to absent contractile gene expression and expression of chondrocyte-like transcripts (e.g., *Sox9, Chad, Col2a1*).^21^ Selected marker genes for the 5 clusters are displayed in **Fig. 4B** and full marker sets included in **Table S3**.

**Figure 4.**
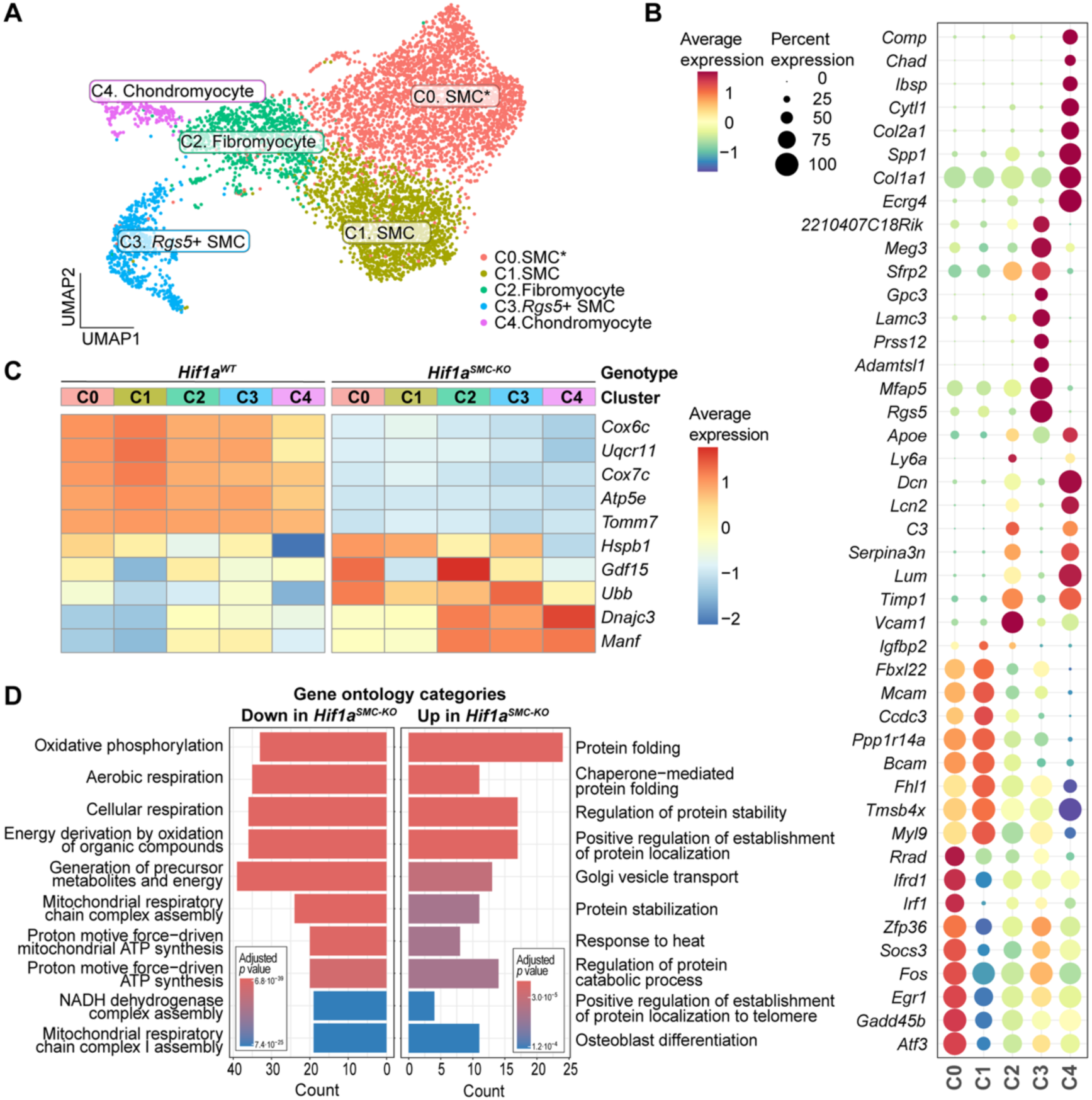
Single-cell RNA-seq reveals changes in OXPHOS and proteostasis genes. **A,** Integrated UMAP of SMCs and SMC-derived cells in atherosclerotic aortic arches from *Hif1a*^WT^ and *Hif1a*^SMC-KO^ mice. Clusters were annotated as C0.SMC* (contractile SMCs with stress-related gene expression), C1.SMC (conventional contractile SMCs), C2.Fibromyocytes (fibroblast-like SMC-derived cells), C3.*Rgs5*+SMC (Rgs5+-expressing SMCs), and C4.Chondromyocytes (chondrocyte-like SMC-derived cells). **B,** Marker genes used to guide the annotation. **C,** Selected genes within top 10 DEGs for each SMC-derived cell clusters, highlighting broad downregulation of OXPHOS/mitochondrial genes (*Cox6c*, *Uqcr11*, *Cox7c*, *Atp5e*, *Tomm7*) and increased expression of protein misfolding and stress response genes (e.g., *Ubb*, *Manf*, *Hspa8*, *Hspb1*, *Dnajc3*) in *Hif1a*^SMC-KO^ versus *Hif1a*^WT^ plaques. **D,** Gene Ontology Biological Process enrichment analysis of up- and downregulated genes in *Hif1a*^SMC-KO^ versus *Hif1a*^WT^ SMC-derived cells. Analysis was performed on pools of aortic arches from 6 *Hif1a*^WT^ and 8 *Hif1a*^SMC-KO^ mice.

We first assessed the expression of known HIF1-regulated genes with a role in glycolysis (*Slc2a1, Ldha, Pdk1*), angiogenesis (*Vegfa*), iron metabolism (*Tfrc*), autophagy (*Bnip3*), and mitochondrial adaptation to hypoxia (*Ndufa4l2, Cox4i2*). Expression of these target genes was enriched in fibromyocytes and chondromyocytes of *Hif1a*^WT^ samples but largely absent in *Hif1a*^SMC-KO^ cells (**Fig. S2A**). A complementary HIF1α gene module score, calculated using a curated list of direct HIF1 targets from MSigDB,^24^ confirmed the reduction in HIF1-dependent transcription in fibromyocytes and chondromyocytes of *Hif1a*^SMC-KO^ mice (**Fig. S2B**).

To investigate other changes resulting from the loss of *Hif1a* expression, we then performed an unbiased analysis of differentially expressed genes (DEGs) in each cluster (**Table S4**). Across SMC-derived clusters in *Hif1a*^SMC-KO^ compared with *Hif1a*^WT^ mice, we observed a coordinated downregulation of mitochondrial and electron transport chain components (e.g., *Cox6c*, *Uqcr11*, *Cox7c*, *Atp5e*, *Tomm7*). Conversely, genes related to protein misfolding and stress response (e.g., *Hspb1*, *Gdf15*, *Ubb*, *Dnajc3, Manf*) were among the most robustly upregulated genes (**Fig. 4C**). Gene ontology analysis of the top upregulated and downregulated genes confirmed these two themes (**Fig. 4D**). Combined, the expression changes suggest that SMC-derived cells in *Hif1a*^SMC-KO^ mice downregulate oxidative phosphorylation and mount compensatory responses to the accumulation of misfolded or damaged protein.

### Hypoxic Hif1a-deficient SMCs accumulate dysfunctional, ROS-producing mitochondria

The observed downregulation of oxidative phosphorylation gene expression in SMC-derived cells from *Hif1a*^SMC-KO^ plaques may seem surprising when considering that a key adaptation to hypoxia is a shift from oxidative phosphorylation to anaerobic glycolysis^25,26^ - loss of HIF1α would be expected to impair this metabolic shift. Given the concomitant signs of cellular stress, however, we reasoned that the downregulation may represent a compensatory mechanism aimed at reducing ROS production, as complexes I and III are major sources of mitochondrial ROS generation, especially under hypoxia .^27^

To evaluate this hypothesis, we conducted a series of *in vitro* studies in immortalized rat aortic SMCs subjected to hypoxia (1% O_2_) combined with *Hif1a* knockdown. SMC cultures under hypoxia showed markedly reduced growth after 48 hours, and this timepoint was selected for subsequent experiments (**Fig. 5A**). *Hif1a* silencing was efficient (≈80% decline in *Hif1a* mRNA expression) and functional, as shown by the attenuation of the hypoxia-induced HIF1 target gene *Slc2a1* (encoding GLUT1) (**Fig. 5B**). Notably, *Hif1a* knockdown increased cell counts under hypoxia (**Fig. 5C**), consistent with the observed expansion of SMC-derived cells in *Hif1a*^SMC-KO^ plaques (**Fig. 2B**). To investigate whether *Hif1a* loss impacts mitochondrial homeostasis during hypoxia, we assessed mitochondrial content and membrane potential using MitoTracker Green and MitoTracker Deep Red, respectively.^28,29^ After 48 hours of hypoxia, subsets of SMCs accumulated depolarized mitochondria (MitoTracker Deep Red ^low^ / MitoTracker Green ^high^) (**Fig. 5D**), *Hif1a* silencing further increased the fraction of SMCs with depolarized mitochondria (**Fig. 5D**). These findings indicate that the loss of *Hif1a* exacerbates the mitochondrial membrane potential disruption triggered by hypoxia in SMCs, pointing towards a protective role of *Hif1a* against mitochondrial damage during low oxygen conditions.

**Figure 5.**
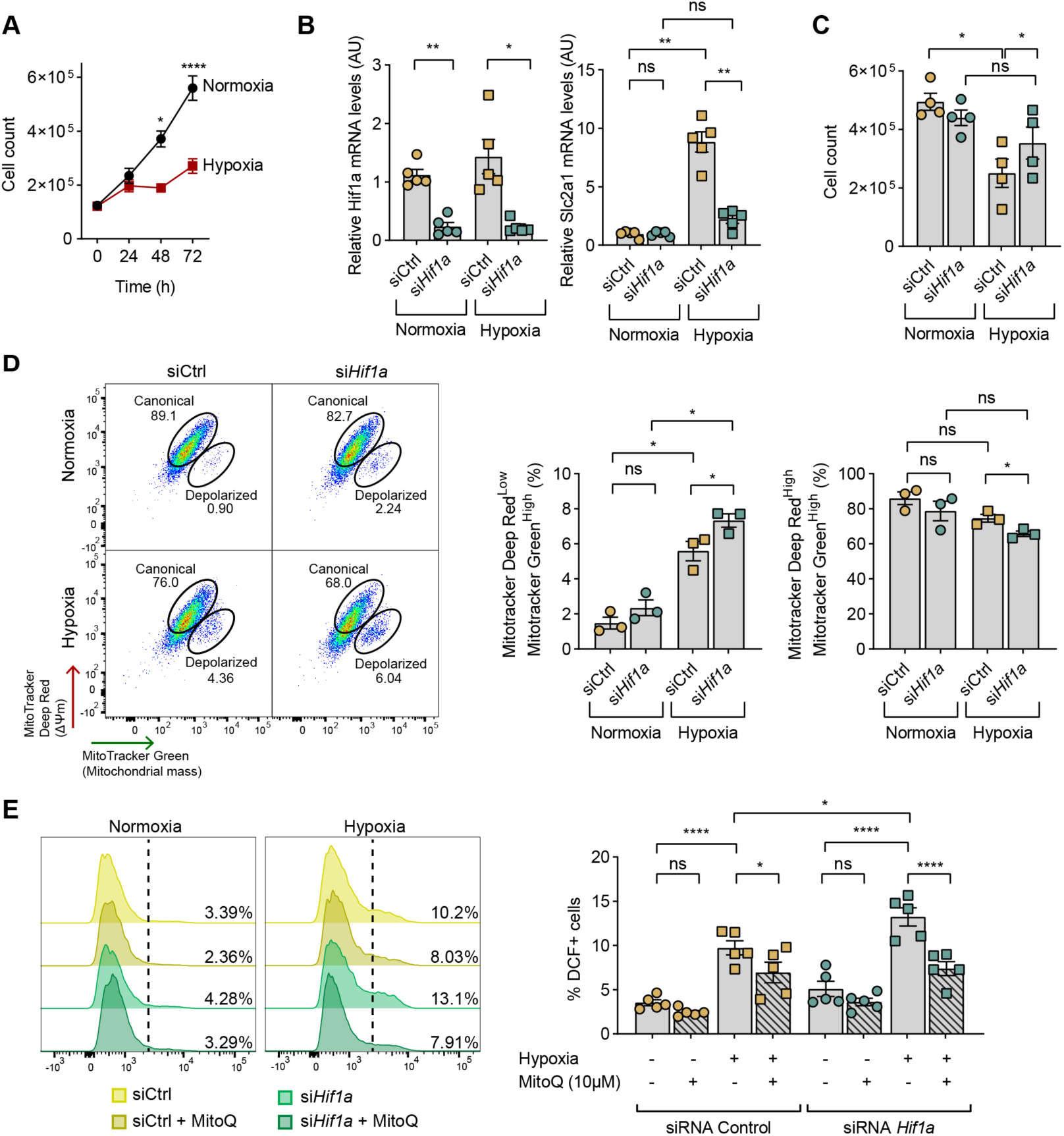
*Hif1a* silencing exacerbates hypoxia-induced impairment of SMC growth, mitochondrial membrane potential, and ROS homeostasis. **A,** Growth curves of immortalized rat SMCs cultured under normoxia or hypoxia (1% O_2_). Cell numbers were significantly reduced by hypoxia at 48 h, the time point chosen for subsequent experiments. **B**, RT-qPCR analysis of *Hif1a* and *Slc2a1* (GLUT1) mRNA after 48 h of normoxia or hypoxia in SMCs transfected with *Hif1a*-targeting siRNA (si*Hif1a*) or a non-targeting control siRNA (si*Ctrl*). *Ywhaz* and *Hprt1* served as reference genes. **C,** Cell counts after 48 h of normoxia or hypoxia in si*Ctrl*- and si*Hif1a*-treated SMCs, showing that *Hif1a* silencing attenuates the hypoxia-induced reduction in cell number. **D,** Flow cytometry of si*Ctrl*- and si*Hif1a*-treated SMCs after 48 h of normoxia or hypoxia. Cells were stained with MitoTracker Green (mitochondrial mass) and MitoTracker Deep Red (membrane potential-sensitive) to distinguish canonical mitochondria (Green^high^/DeepRed^high^) from depolarized mitochondria (Green^high^/DeepRed^low^); DAPI was used to exclude dead cells. **E,** ROS production assessed by DCF fluorescence (H_2_DCFDA) measured by flow cytometry, with or without the mitochondria-targeted antioxidant mitoquinone (MitoQ), confirming mitochondrial origin of the hypoxia- and si*Hif1a*-induced ROS signal. Bars, mean ± SEM. Data points represent the average of multiple wells in an independent experiment, with experiments conducted 3 (D), 4 (A-C) or 5 (E) times. Groups were compared with two-way ANOVA with Sidak’s post hoc test (A), repeated-measures one-way ANOVA (B-D) or two-way ANOVA (E) with Tukey’s post hoc test. \**P*<0.05, \*\**P*<0.01, \*\*\*\**P*<0.0001.

Loss of membrane potential has previously been associated with ROS-generating mitochondria.^29^ To measure ROS production in cultured SMCs, we used the H_2_DCFDA probe, which, once internalized, undergoes deacetylation by intracellular esterases to H_2_DCF. Electron loss generates the highly fluorescent 2’,7’-dichlorofluorescein (DCF), which is proportional to general cellular oxidation.^30^ We found that si*Hif1a*-transfected cells produce higher levels of ROS, indicating an elevated state of oxidative stress. Because H₂DCFDA reflects global ROS production and lacks mitochondrial specificity, we co-incubated cultures with the mitochondria-targeted antioxidant mitoquinone (MitoQ).^31^ Treatment with MitoQ alleviated the DCF⁺ signal in both control and *siHif1a* samples, indicating that the rise in the oxidative signal during hypoxia and the amplifying effect of *Hif1a* silencing originates from increased mitochondrial oxidant species (**Fig. 5E**).

Together, these *in vitro* results show that the loss of *Hif1a* in SMCs affects their mitochondrial function resulting in increased ROS production, letting us to suggest that the downregulation of mitochondrial respiratory/OXPHOS genes observed in the single-cell RNA-seq data may be a compensatory response to sustained ROS overproduction in response to *Hif1a* deletion.

### Oxidative and ER stress markers accumulate in SMC-derived cells of Hif1a^SMC-KO^ plaques

Given the *in vitro* evidence that HIF1α loss predisposes SMCs to oxidative stress under hypoxic conditions, we next asked whether this was reflected in the gene expression of SMC-derived cells from *Hif1a*^SMC-KO^ plaques. Single-cell gene-set scoring of the scRNA-seq dataset revealed enrichment of the “cellular response to oxidative stress” gene module (GO:0034599) in *Hif1a*^SMC-KO^ plaques relative to controls (**Fig. 6A**). To validate this transcriptomic signal histologically, we assessed oxidative DNA damage within the SMC lineage by staining for 8-hydroxy-2′-deoxyguanosine (8OHdG), which revealed a higher abundance of tdT^+^8OHdG^+^ SMC-derived cells in *Hif1a*^SMC-KO^ plaques (**Fig. 6B-C**). Notably, 8OHdG accumulated in SOX9^+^ chondromyocyte-rich regions, prompting quantification of triple-positive cells (tdT^+^SOX9^+^8OHdG^+^). This analysis confirmed that chondromyocytes - the SMC-derived subpopulation with stronger upregulation of hypoxia-related genes under wild-type conditions (**Fig. 4C**) - are vulnerable to oxidative stress in the absence of HIF1α (**Fig. 6B-C**).

**Figure 6.**
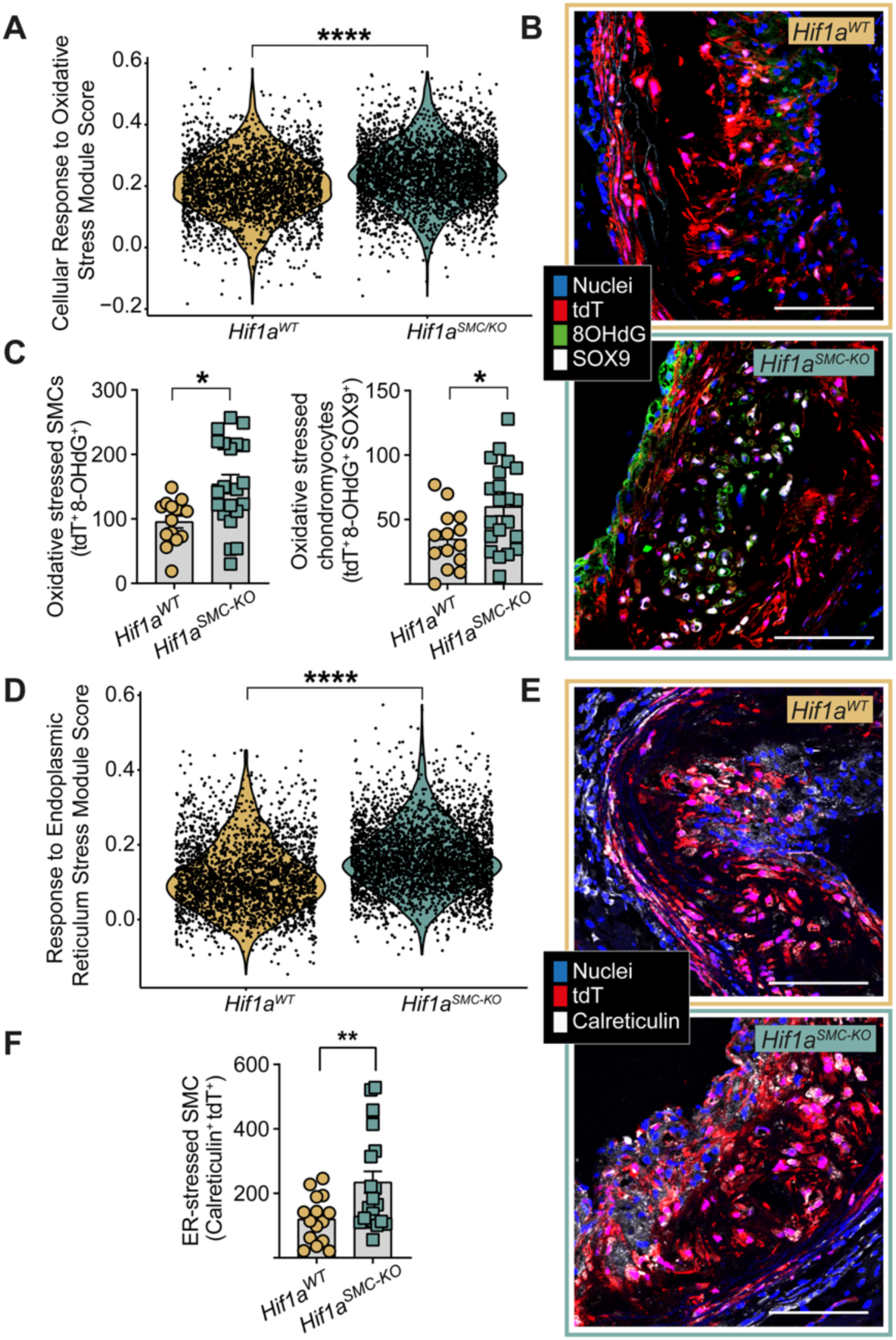
*Hif1a* deletion in SMC-derived plaque cells increases markers of oxidative damage and proteostatic stress. **A,** Expression of the “cellular response to oxidative stress” gene module (GO:0034599) in plaque SMC-derived cells of the scRNA-seq data set, showing upregulation in *Hif1a*^SMC-KO^ compared with *Hif1a*^WT^ mice. **B,** Representative immunostaining of aortic root sections for 8OHdG (oxidative DNA damage) and SOX9 (chondromyocytes). TdT marks SMC-derived cells. **C**, Quantification of 8OHdG⁺ tdT⁺ and 8OHdG⁺ tdT⁺ SOX9⁺ cells, showing increase of both in *Hif1a*^SMC-KO^ plaques. **D,** Expression of the “response to endoplasmic reticulum stress” gene module (GO:0034976) in plaque SMC-derived cells of the scRNA-seq data set, showing upregulation in *Hif1a*^SMC-KO^ compared with *Hif1a*^WT^ mice. **E,** Representative immunostaining for calreticulin (an ER chaperone) in aortic root plaque. **F,** Quantification of Calreticulin⁺ tdT⁺, showing an increase in *Hif1a*^SMC-KO^ samples. Bars, mean ± SEM. Scale bars, 100µm. Data were obtained from 14-15 *Hif1a^WT^* and 21 *Hif1a^SMC-KO^* mice, and groups were compared by unpaired t-tests. \**P*<0.05, \*\**P*<0.01, \*\*\*\**P*<0.0001.

The close mechanistic links between oxidative and proteostatic stress, and the upregulation of several protein misfolding genes in scRNA-seq data (**Fig. 4C**), prompted us to examine whether the unfolded protein response was elevated in plaque SMC-derived cells of *Hif1a*^SMC-KO^ mice. ScRNA-seq module scoring confirmed enrichment of the “response to endoplasmic reticulum stress” gene module (GO:0034976) in *Hif1a*^SMC-KO^ plaque SMC-derived cells (**Fig. 6D**). To corroborate this histologically, we stained for calreticulin, an ER chaperone upregulated during conditions of increased protein folding demand, which was increased in *Hif1a*^SMC-KO^ lesions and enriched within the SMC-derived compartment (**Fig. 6E-F**).

Altogether, these findings point to a coordinated increase in oxidative damage and protein misfolding in SMC-derived cells of *Hif1a*^SMC-KO^ plaques, suggesting that loss of HIF1α-mediated adaptation to hypoxia renders SMC-derived cells vulnerable to stress pathways that may contribute to plaque expansion and increased necrotic core formation.

## Discussion

In the present study, we demonstrate that loss of HIF1α signaling in SMC-derived cells leads to larger plaques with more necrosis and calcification, accompanied by accelerated accumulation of all SMC-derived cell types – particularly chondrocyte-like cells – and elevated markers of oxidative stress, cell proliferation, and apoptosis. Together, these findings identify HIF1α as a protective regulator of SMC-derived cells in advanced atherosclerosis.

Hypoxia develops in advanced atherosclerotic plaques when oxygen demand exceeds supply,^32^ as demonstrated in both mouse and human lesions using oxygen sensing probes.^33,34^ SMC-derived fibromyocytes and chondromyocytes accumulate selectively in the hypoxic plaque interior, yet how these cells respond to low oxygen – and how that response shapes their accumulation and fate – have remained poorly understood. One prior study, by Liu et al., found that SM22α-Cre-induced *Hif1a* deletion attenuated vascular lipid accumulation and inflammation in pressure-accelerated atherosclerosis.^35^ Although this appears to conflict with our observations, key differences in the examined disease stage likely reconcile them: Liu et al. examined early, inflammation-dominated lesions in which SMC-derived cell expansion plays little role, whereas we studied later-stage atherogenesis where SMC-derived cells dominate the plaque.^35^ Such stage-dependent duality is not unusual in atherosclerosis; IL-1 isoforms, for example, exert opposing effects in early versus late atherosclerotic lesions.^36^

Although HIF1α classically promotes glycolysis and represses mitochondrial respiration via PDK1 induction,^37^ its loss in SMC-derived plaque cells did not trigger compensatory upregulation of oxidative metabolism genes. Rather, the scRNA-seq analysis and staining data revealed mitochondrial dysfunction and proteostatic stress as the dominant signatures of *Hif1a* deletion. Cell culture studies supported the underlying mechanistic links by showing that *Hif1a* silencing increased cell counts, aggravated mitochondrial depolarization, and elevated intracellular ROS — an effect abolished by the mitochondria-targeted antioxidant MitoQ, confirming a mitochondrial origin. Taken together, these findings indicate that a main function of HIF1α in hypoxic SMC-derived cells is to safeguard against the mitochondrial ROS production that results from partial electron transport chain blockade caused by low oxygen. Uncontrolled mitochondrial ROS production can drive proliferation of vascular SMCs,^38,39^ while oxidative protein damage directly triggers proteostatic stress and unfolded protein response activation.^40^ As the unfolded protein response has been shown to promote SMC dedifferentiation and plaque growth in vivo,^41^ both pathways provide credible mechanisms for the expanded SMC-derived compartment and enlarged necrotic core observed in *Hif1a*^SMC-KO^ plaques.

The expansion of SMC-derived cells in *Hif1a*^SMC-KO^ plaques was particularly prominent among *Col2a1*-expressing chondromyocytes, suggesting that this subpopulation is particularly reliant on HIF1α for adaptation to the hypoxic plaque environment. This parallels developmental studies of cartilage formation in the hypoxic fetal growth plate, where *Hif1a* loss in developing chondrocytes causes severe cell death and disordered differentiation with loss of growth arrest.^42,43^ Notably, genetic suppression of mitochondrial respiration rescued this phenotype,^44^ implicating excessive mitochondrial oxygen consumption — and ROS generation — as the proximate cause of cell death in the absence of HIF1α, a mechanism likely operative in plaque chondromyocytes as well.

Our study adds to the complex, cell type-specific picture of HIF1 signaling in atherosclerosis. In macrophages, HIF1α exacerbates disease by promoting inflammation and foam cell formation,^45^ whereas in endothelial cells it facilitates neovascularization and monocyte recruitment,^6,46^ and in dendritic cells and T cells, by contrast, it exerts a protective effect.^47,48^ The opposing effects of HIF1 signaling across different cell types make it difficult to predict the net outcome of global HIF1 inhibition. This is of particular relevance to HIF1 inhibitors which are under investigation as anticancer agents,^49,50^ exploiting the tumor dependence on hypoxia-driven metabolism. Given that cardiovascular disease is highly prevalent in cancer patients, our results raise the possibility that systemic HIF1 inhibition could have unintended consequences in the vasculature – accelerating oxidative stress, SMC-derived cell death, and necrotic core expansion. This warrants consideration in the safety monitoring of HIF-targeting therapies, particularly in patients with known or suspected advanced atherosclerosis.

Our study has several limitations. First, the proposed mechanisms involving mitochondrial ROS were inferred from gene expression analysis and cell culture experiments rather than direct functional measurements in vivo. Future studies with selective targeting of mitochondrial respiration or antioxidant defenses specifically in SMC-derived cells will be important to establish causality. Second, our findings were obtained in a mouse model of atherosclerosis, which differs from human disease in important ways. Most notably, advanced human plaques typically develop extensive microvascular networks, which may partially relieve hypoxia and reduce SMC dependence on HIF1-driven adaptation. Whether the protective role of HIF1α in SMC-derived cells extends to human atheromas – particularly in well-perfused regions – therefore remains to be established and represents an important direction for future investigation.

## Nonstandard Abbreviations and Acronyms

AAV: Adeno-Associated Virus
DCFDA: Dichlorodihydrofluorescein diacetate
ER/ER-stress: Endoplasmic-Reticulum/Endoplasmid Reticulum stress
ETC: Electron Transport Chain
HFD: High-Fat Diet
HIF: Hypoxia-Inducible Factor
HRE: Hypoxia-Response Element
ISH: In Situ Hybridization
MYH11/Myh11: Myosin Heave Chain 11
OCR: Oxygen Consumption Rate
OXPHOS: Oxidative Phosphorylation
PCSK9: Proprotein Convertase Subtilisin/Kexin type 9
ROS: Reactive Oxygen Species
SMC: Smooth Muscle Cell
tdT: tdTomato
VHL: Von Hippel-Lindau

## Acknowledgments

We thank Leticia González-Cintado and the members of the CNIC Viral Vectors Unit, Microscopy Unit, Genomics Unit, Bioinformatics Unit, Cellomics Unit and Animal Facility for excellent technical help.

## References

1. Sluimer JC, Gasc JM, van Wanroij JL, Kisters N, Groeneweg M, Sollewijn Gelpke MD, Cleutjens JP, van den Akker LH, Corvol P, Wouters BG, Daemen MJ, Bijnens APJ. Hypoxia, Hypoxia-Inducible Transcription Factor, and Macrophages in Human Atherosclerotic Plaques Are Correlated With Intraplaque Angiogenesis. J Am Coll Cardiol. 2008;51(13):1258–1265. doi:10.1016/j.jacc.2007.12.025

2. Al-Mashhadi RH, Tolbod LP, Bloch LØ, Bjørklund MM, Nasr ZP, Al-Mashhadi Z, Winterdahl M, Frøkiær J, Falk E, Bentzon JF. 18Fluorodeoxyglucose Accumulation in Arterial Tissues Determined by PET Signal Analysis. J Am Coll Cardiol. 2019;74(9):1220–1232. doi:10.1016/j.jacc.2019.06.057

3. Parathath S, Mick SL, Feig JE, Joaquin V, Grauer L, Habiel DM, Gassmann M, Gardner LB, Fisher EA. Hypoxia is present in murine atherosclerotic plaques and has multiple adverse effects on macrophage lipid metabolism. Circ Res. 2011;109(10):1141–1152. doi:10.1161/CIRCRESAHA.111.246363

4. Crawford DW, Back LH, Cole MA. In vivo oxygen transport in the normal rabbit femoral arterial wall. J Clin Invest. 1980;65(6):1498–1508. doi:10.1172/JCI109815

5. Marsch E, Sluimer JC, Daemen MJAP. Hypoxia in atherosclerosis and inflammation. Curr Opin Lipidol. 2013;24(5):393–400. doi:10.1097/MOL.0b013e32836484a4

6. Akhtar S, Hartmann P, Karshovska E, Rinderknecht FA, Subramanian P, Gremse F, Grommes J, Jacobs M, Kiessling F, Weber C, Steffens S, Schober A. Endothelial Hypoxia-Inducible Factor-1α Promotes Atherosclerosis and Monocyte Recruitment by Upregulating MicroRNA-19a. Hypertension. 2015;66(6):1220–1226. doi:10.1161/HYPERTENSIONAHA.115.05886

7. Basatemur GL, Jørgensen HF, Clarke MCH, Bennett MR, Mallat Z. Vascular smooth muscle cells in atherosclerosis. Nat Rev Cardiol. 2019;16(12):727–744. doi:10.1038/s41569-019-0227-9

8. Owens GK, Kumar MS, Wamhoff BR. Molecular regulation of vascular smooth muscle cell differentiation in development and disease. Physiol Rev. 2004;84(3):767–801. doi:10.1152/physrev.00041.2003

9. Jacobsen K, Lund MB, Shim J, Gunnersen S, Füchtbauer EM, Kjolby M, Carramolino L, Bentzon JF. Diverse cellular architecture of atherosclerotic plaque derives from clonal expansion of a few medial SMCs. JCI Insight. 2017;2(19):e95890, 95890. doi:10.1172/jci.insight.95890

10. Chen R, McVey DG, Shen D, Huang X, Ye S. Phenotypic Switching of Vascular Smooth Muscle Cells in Atherosclerosis. J Am Heart Assoc. 2023;12(20):e031121. doi:10.1161/JAHA.123.031121

11. Kaelin WG, Ratcliffe PJ. Oxygen sensing by metazoans: the central role of the HIF hydroxylase pathway. Mol Cell. 2008;30(4):393–402. doi:10.1016/j.molcel.2008.04.009

12. Cash TP, Pan Y, Simon MC. Reactive oxygen species and cellular oxygen sensing. Free Radic Biol Med. 2007;43(9):1219–1225. doi:10.1016/j.freeradbiomed.2007.07.001

13. Hamanaka RB, Chandel NS. Mitochondrial reactive oxygen species regulate hypoxic signaling. Curr Opin Cell Biol. 2009;21(6):894–899. doi:10.1016/j.ceb.2009.08.005

14. Guzy RD, Schumacker PT. Oxygen sensing by mitochondria at complex III: the paradox of increased reactive oxygen species during hypoxia. Exp Physiol. 2006;91(5):807–819. doi:10.1113/expphysiol.2006.033506

15. Chandel NS, McClintock DS, Feliciano CE, Wood TM, Melendez JA, Rodriguez AM, Schumacker PT. Reactive oxygen species generated at mitochondrial complex III stabilize hypoxia-inducible factor-1alpha during hypoxia: a mechanism of O2 sensing. J Biol Chem. 2000;275(33):25130–25138. doi:10.1074/jbc.M001914200

16. Li HS, Zhou YN, Li L, Li SF, Long D, Chen XL, Zhang JB, Feng L, Li YP. HIF-1α protects against oxidative stress by directly targeting mitochondria. Redox Biol. 2019;25:101109. doi:10.1016/j.redox.2019.101109

17. Papandreou I, Cairns RA, Fontana L, Lim AL, Denko NC. HIF-1 mediates adaptation to hypoxia by actively downregulating mitochondrial oxygen consumption. Cell Metab. 2006;3(3):187–197. doi:10.1016/j.cmet.2006.01.012

18. Ryan HE, Poloni M, McNulty W, Elson D, Gassmann M, Arbeit JM, Johnson RS. Hypoxia-inducible factor-1alpha is a positive factor in solid tumor growth. Cancer Res. 2000;60(15):4010–4015.

19. Hamidi A, Wolf A, Dueva R, Kaufmann M, Göpelt K, Iliakis G, Metzen E. Depletion of HIF-1α by Inducible Cre/loxP Increases the Sensitivity of Cultured Murine Hepatocytes to Ionizing Radiation in Hypoxia. Cells. 2022;11(10):1671. doi:10.3390/cells11101671

20. Carramolino L, Albarrán-Juárez J, Markov A, Hernández-SanMiguel E, Sharysh D, Cumbicus V, Morales-Cano D, Labrador-Cantarero V, Møller PL, Nogales P, Benguria A, Dopazo A, Sanchez-Cabo F, Torroja C, Bentzon JF. Cholesterol lowering depletes atherosclerotic lesions of smooth muscle cell-derived fibromyocytes and chondromyocytes. Nat Cardiovasc Res. 2024;3(2):203–220. doi:10.1038/s44161-023-00412-w

21. Kim JB, Zhao Q, Nguyen T, Pjanic M, Cheng P, Wirka R, Travisano S, Nagao M, Kundu R, Quertermous T. Environment-Sensing Aryl Hydrocarbon Receptor Inhibits the Chondrogenic Fate of Modulated Smooth Muscle Cells in Atherosclerotic Lesions. Circulation. 2020;142(6):575–590. doi:10.1161/CIRCULATIONAHA.120.045981

22. van den Brink SC, Sage F, Vértesy Á, Spanjaard B, Peterson-Maduro J, Baron CS, Robin C, van Oudenaarden A. Single-cell sequencing reveals dissociation-induced gene expression in tissue subpopulations. Nat Methods. 2017;14(10):935–936. doi:10.1038/nmeth.4437

23. Wirka RC, Wagh D, Paik DT, Pjanic M, Nguyen T, Miller CL, Kundu R, Nagao M, Coller J, Koyano TK, Fong R, Woo YJ, Liu B, Montgomery SB, Wu JC, Zhu K, Chang R, Alamprese M, Tallquist MD, Kim JB, Quertermous T. Atheroprotective roles of smooth muscle cell phenotypic modulation and the TCF21 disease gene as revealed by single-cell analysis. Nat Med. 2019;25(8):1280–1289. doi:10.1038/s41591-019-0512-5

24. Semenza GL. Hypoxia-inducible factor 1: oxygen homeostasis and disease pathophysiology. Trends Mol Med. 2001;7(8):345–350. doi:10.1016/s1471-4914(01)02090-1

25. Gleadle JM, Ratcliffe PJ. Induction of hypoxia-inducible factor-1, erythropoietin, vascular endothelial growth factor, and glucose transporter-1 by hypoxia: evidence against a regulatory role for Src kinase. Blood. 1997;89(2):503–509.

26. Airley R, Loncaster J, Davidson S, Bromley M, Roberts S, Patterson A, Hunter R, Stratford I, West C. Glucose transporter glut-1 expression correlates with tumor hypoxia and predicts metastasis-free survival in advanced carcinoma of the cervix. Clin Cancer Res Off J Am Assoc Cancer Res. 2001;7(4):928–934.

27. Balogh E, Tóth A, Méhes G, Trencsényi G, Paragh G, Jeney V. Hypoxia Triggers Osteochondrogenic Differentiation of Vascular Smooth Muscle Cells in an HIF-1 (Hypoxia-Inducible Factor 1)–Dependent and Reactive Oxygen Species–Dependent Manner. Arterioscler Thromb Vasc Biol. 2019;39(6):1088–1099. doi:10.1161/ATVBAHA.119.312509

28. Greene AW, Grenier K, Aguileta MA, Muise S, Farazifard R, Haque ME, McBride HM, Park DS, Fon EA. Mitochondrial processing peptidase regulates PINK1 processing, import and Parkin recruitment. EMBO Rep. 2012;13(4):378–385. doi:10.1038/embor.2012.14

29. Zhou R, Yazdi AS, Menu P, Tschopp J. A role for mitochondria in NLRP3 inflammasome activation. Nature. 2011;469(7329):221–225. doi:10.1038/nature09663

30. Eruslanov E, Kusmartsev S. Identification of ROS using oxidized DCFDA and flow-cytometry. Methods Mol Biol. 2010;594:57–72. doi:10.1007/978-1-60761-411-1_4

31. Bennett NK, Lee M, Orr AL, Nakamura K. Systems-level analyses dissociate genetic regulators of reactive oxygen species and energy production. Proc Natl Acad Sci. 2024;121(3):e2307904121. doi:10.1073/pnas.2307904121

32. Björnheden T, Levin M, Evaldsson M, Wiklund O. Evidence of Hypoxic Areas Within the Arterial Wall In Vivo. Arterioscler Thromb Vasc Biol. 1999;19(4):870–876. doi:10.1161/01.ATV.19.4.870

33. Van Herck JL, De Meyer GRY, Martinet W, Van Hove CE, Foubert K, Theunis MH, Apers S, Bult H, Vrints CJ, Herman AG. Impaired Fibrillin-1 Function Promotes Features of Plaque Instability in Apolipoprotein E–Deficient Mice. Circulation. 2009;120(24):2478–2487. doi:10.1161/CIRCULATIONAHA.109.872663

34. Sluimer JC, Gasc JM, van Wanroij JL, Kisters N, Groeneweg M, Sollewijn Gelpke MD, Cleutjens JP, van den Akker LH, Corvol P, Wouters BG, Daemen MJ, Bijnens APJ. Hypoxia, hypoxia-inducible transcription factor, and macrophages in human atherosclerotic plaques are correlated with intraplaque angiogenesis. J Am Coll Cardiol. 2008;51(13):1258–1265. doi:10.1016/j.jacc.2007.12.025

35. Liu D, Lei L, Desir M, Huang Y, Cleman J, Jiang W, Fernandez-Hernando C, Di Lorenzo A, Sessa WC, Giordano FJ. Smooth Muscle Hypoxia-Inducible Factor 1α Links Intravascular Pressure and Atherosclerosis—Brief Report. Arterioscler Thromb Vasc Biol. 2016;36(3):442–445. doi:10.1161/ATVBAHA.115.306861

36. Vromman A, Ruvkun V, Shvartz E, Wojtkiewicz G, Santos Masson G, Tesmenitsky Y, Folco E, Gram H, Nahrendorf M, Swirski FK, Sukhova GK, Libby P. Stage-dependent differential effects of interleukin-1 isoforms on experimental atherosclerosis. Eur Heart J. 2019;40(30):2482–2491. doi:10.1093/eurheartj/ehz008

37. Kim J whan, Tchernyshyov I, Semenza GL, Dang CV. HIF-1-mediated expression of pyruvate dehydrogenase kinase: a metabolic switch required for cellular adaptation to hypoxia. Cell Metab. 2006;3(3):177–185. doi:10.1016/j.cmet.2006.02.002

38. Madamanchi NR, Moon SK, Hakim ZS, Clark S, Mehrizi A, Patterson C, Runge MS. Differential activation of mitogenic signaling pathways in aortic smooth muscle cells deficient in superoxide dismutase isoforms. Arterioscler Thromb Vasc Biol. 2005;25(5):950–956. doi:10.1161/01.ATV.0000161050.77646.68

39. Glogowski PA, Fogacci F, Algieri C, Cugliari A, Trombetti F, Nesci S, Cicero AFG. Reprogramming the Mitochondrion in Atherosclerosis: Targets for Vascular Protection. Antioxidants. 2025;14(12):1462. doi:10.3390/antiox14121462

40. Poletto M, Yang D, Fletcher SC, Vendrell I, Fischer R, Legrand AJ, Dianov GL. Modulation of proteostasis counteracts oxidative stress and affects DNA base excision repair capacity in ATM-deficient cells. Nucleic Acids Res. 2017;45(17):10042–10055. doi:10.1093/nar/gkx635

41. Chattopadhyay A, Kwartler CS, Kaw K, Li Y, Kaw A, Chen J, LeMaire SA, Shen YH, Milewicz DM. Cholesterol-Induced Phenotypic Modulation of Smooth Muscle Cells to Macrophage/Fibroblast-like Cells Is Driven by an Unfolded Protein Response. Arterioscler Thromb Vasc Biol. 2021;41(1):302–316. doi:10.1161/ATVBAHA.120.315164

42. Schipani E, Ryan HE, Didrickson S, Kobayashi T, Knight M, Johnson RS. Hypoxia in cartilage: HIF-1α is essential for chondrocyte growth arrest and survival. Genes Dev. 2001;15(21):2865–2876. doi:10.1101/gad.934301

43. Pfander D, Kobayashi T, Knight MC, Zelzer E, Chan DA, Olsen BR, Giaccia AJ, Johnson RS, Haase VH, Schipani E. Deletion of *Vhlh* in chondrocytes reduces cell proliferation and increases matrix deposition during growth plate development. Development. 2004;131(10):2497–2508. doi:10.1242/dev.01138

44. Yao Q, Khan MP, Merceron C, LaGory EL, Tata Z, Mangiavini L, Hu J, Vemulapalli K, Chandel NS, Giaccia AJ, Schipani E. Suppressing Mitochondrial Respiration Is Critical for Hypoxia Tolerance in the Fetal Growth Plate. Dev Cell. 2019;49(5):748–763.e7. doi:10.1016/j.devcel.2019.04.029

45. Aarup A, Pedersen TX, Junker N, Christoffersen C, Bartels ED, Madsen M, Nielsen CH, Nielsen LB. Hypoxia-Inducible Factor-1α Expression in Macrophages Promotes Development of Atherosclerosis. Arterioscler Thromb Vasc Biol. 2016;36(9):1782–1790. doi:10.1161/ATVBAHA.116.307830

46. Zimna A, Kurpisz M. Hypoxia-Inducible Factor-1 in Physiological and Pathophysiological Angiogenesis: Applications and Therapies. BioMed Res Int. 2015;2015:549412. doi:10.1155/2015/549412

47. Chaudhari SM, Sluimer JC, Koch M, Theelen TL, Manthey HD, Busch M, Caballero-Franco C, Vogel F, Cochain C, Pelisek J, Daemen MJ, Lutz MB, Görlach A, Kissler S, Hermanns HM, Zernecke A. Deficiency of HIF1α in Antigen-Presenting Cells Aggravates Atherosclerosis and Type 1 T-Helper Cell Responses in Mice. Arterioscler Thromb Vasc Biol. 2015;35(11):2316–2325. doi:10.1161/ATVBAHA.115.306171

48. Ben-Shoshan J, Afek A, Maysel-Auslender S, Barzelay A, Rubinstein A, Keren G, George J. HIF-1alpha overexpression and experimental murine atherosclerosis. Arterioscler Thromb Vasc Biol. 2009;29(5):665–670. doi:10.1161/ATVBAHA.108.183319

49. Zhe N, Chen S, Zhou Z, Liu P, Lin X, Yu M, Cheng B, Zhang Y, Wang J. HIF-1α inhibition by 2-methoxyestradiol induces cell death via activation of the mitochondrial apoptotic pathway in acute myeloid leukemia. Cancer Biol Ther. 2016;17(6):625–634. doi:10.1080/15384047.2016.1177679

50. Jin J, Qiu S, Wang P, Liang X, Huang F, Wu H, Zhang B, Zhang W, Tian X, Xu R, Shi H, Wu X. Cardamonin inhibits breast cancer growth by repressing HIF-1α-dependent metabolic reprogramming. J Exp Clin Cancer Res CR. 2019;38(1):377. doi:10.1186/s13046-019-1351-4

51. Bankhead P, Loughrey MB, Fernández JA, Dombrowski Y, McArt DG, Dunne PD, McQuaid S, Gray RT, Murray LJ, Coleman HG, James JA, Salto-Tellez M, Hamilton PW. QuPath: Open source software for digital pathology image analysis. Sci Rep. 2017;7(1):16878. doi:10.1038/s41598-017-17204-5

52. Elvidge GP, Glenny L, Appelhoff RJ, Ratcliffe PJ, Ragoussis J, Gleadle JM. Concordant regulation of gene expression by hypoxia and 2-oxoglutarate-dependent dioxygenase inhibition: the role of HIF-1alpha, HIF-2alpha, and other pathways. J Biol Chem. 2006;281(22):15215–15226. doi:10.1074/jbc.M511408200

53. Reilly CF. Rat vascular smooth muscle cells immortalized with SV40 large T antigen possess defined smooth muscle cell characteristics including growth inhibition by heparin. J Cell Physiol. 1990;142(2):342–351. doi:10.1002/jcp.1041420217

54. Presley AD, Fuller KM, Arriaga EA. MitoTracker Green labeling of mitochondrial proteins and their subsequent analysis by capillary electrophoresis with laser-induced fluorescence detection. J Chromatogr B Analyt Technol Biomed Life Sci. 2003;793(1):141–150. doi:10.1016/s1570-0232(03)00371-4

55. Agnello M, Morici G, Rinaldi AM. A method for measuring mitochondrial mass and activity. Cytotechnology. 2008;56(3):145–149. doi:10.1007/s10616-008-9143-2

56. Monteiro L de B, Davanzo GG, de Aguiar CF, Moraes-Vieira PMM. Using flow cytometry for mitochondrial assays. MethodsX. 2020;7:100938. doi:10.1016/j.mex.2020.100938

57. Xiao B, Deng X, Zhou W, Tan EK. Flow Cytometry-Based Assessment of Mitophagy Using MitoTracker. Front Cell Neurosci. 2016;10. doi:10.3389/fncel.2016.00076

58. Gomes A, Fernandes E, Lima JLFC. Fluorescence probes used for detection of reactive oxygen species. J Biochem Biophys Methods. 2005;65(2-3):45–80. doi:10.1016/j.jbbm.2005.10.003

59. LeBel CP, Ischiropoulos H, Bondy SC. Evaluation of the probe 2’,7’-dichlorofluorescin as an indicator of reactive oxygen species formation and oxidative stress. Chem Res Toxicol. 1992;5(2):227–231. doi:10.1021/tx00026a012

60. Frühwirth M, Ruedl C, Ellemunter H, Böck G, Wolf H. Flow-cytometric evaluation of oxidative burst in phagocytic cells of children with cystic fibrosis. Int Arch Allergy Immunol. 1998;117(4):270–275. doi:10.1159/000024022

